# RaPID-Query for Fast Identity by Descent Search and Genealogical Analysis

**DOI:** 10.1101/2022.02.03.478907

**Authors:** Yuan Wei, Ardalan Naseri, Degui Zhi, Shaojie Zhang

## Abstract

The size of genetic databases has grown large enough such that, genetic genealogical search, a process of inferring familial relatedness by identifying DNA matches, has become a viable approach to help individuals finding missing family members or law enforcement agencies locating suspects. However, a fast and accurate method is needed to search an out-of-database individual against millions of individuals in such databases. Most existing approaches only offer all-vs-all within panel match. Some prototype algorithms offer 1-vs-all query from out-of-panel individual, but they do not tolerate errors. A new method, random projection-based identical-by-descent (IBD) detection (RaPID) query, referred as RaPID-Query, is introduced to make fast genealogical search possible. RaPID-Query method identifies IBD segments between a query haplotype and a panel of haplotypes. By integrating matches over multiple PBWT indexes, RaPID-Query method is able to locate IBD segments quickly with a given cutoff length while allowing mismatched sites in IBD segments. A single query against all UK biobank autosomal chromosomes can be completed within 2.76 seconds CPU time on average, with the minimum 7 cM IBD segment length and minimum 700 markers. Using the same criteria, RaPID-Query can achieve 0.099 false negative rate and 0.017 false positive rate at the same time on a chromosome 20 sequencing panel having 92,296 sites, which is comparable to the state-of-the-art IBD detection method Hap-IBD. For the relatedness degree separation experiments, RaPID-Query is able to distinguish up to fourth degree of the familial relatedness for a given individual pair, and the area under the receiver operating characteristic curve values are at least 97.28%. It is anticipated that RaPID-Query will make genealogical search convenient and effective, potentially with the integration of complex inference models.

## 1 Introduction

With the popularity of direct-to-consumer (DTC) genotyping services, genetic databases are growing to tens of millions of individuals (https://thednageek.com/dna-tests/). Thus, using single nucleotide polymorphisms (SNPs) data from autosomal chromosomes to perform genealogical search has become feasible for familial relationship inference. DTC companies provide such genealogy service to predict the degree of relationships using accumulated shared identical-by-descent (IBD) segments between each customer and the genetic database of collected genetic data of their customers. In early days, IBD segments are identified from a pairwise comparison based detection algorithm (Henn *et al*., 2012). Currently, there are efficient IBD detection methods emerged and available for relatedness inference, such as PBWT (Durbin, 2014), PBWT-Query (Naseri *et al*., 2019a), RaPID (Naseri *et al*., 2019b), Hap-IBD (Zhou *et al*., 2020a), FastSMC (Nait Saada *et al*., 2020), TPBWT (Freyman *et al*., 2020), d-PBWT (Sanaullah *et al*., 2021), iLASH (Shemirani *et al*., 2021). Though these latest methods show the potential power of IBD driven approach in relatedness studies (Sticca *et al*., 2021), most of them are not able to perform query search, a process that takes an individual’s genetic data as the query and searches the population genetic database for IBD segments.

For real-time query search, indexing-based approach such as Naseri et al.’s and Sanaullah et al.’s query approach (Naseri *et al*., 2019a; Sanaullah *et al*., 2021) are promising. Both query approaches are based on the Positional Burrows–Wheeler transform (PBWT), an efficient haplotype matching algorithm developed by Durbin (2014). Naseri et al. proposes PBWT-Query, a long match based query algorithm that is able to effectively find IBD segments in a given panel for a given query. It is more practical formulation for genealogical search than Durbin’s set-maximal match based query algorithm, as it allows specifying the minimal length of the match. Naseri et al. also proposes L-PBWT-Query which uses an additional data structure LEAP array to achieve run-time efficiency (Naseri *et al*., 2019a). Sanaullah et al. further improves the PBWT-Query algorithm by proposing the triple sweep long match query and the single sweep long match query algorithms. Comparing to Naseri et al.’s algorithm, Sanaullah et al.’s single sweep long match query algorithm efficiently reduces the memory usage while keeps a fast query time. The triple sweep long match query algorithm theoretically achieves the linear run time complexity. For chromosome 21 panel with 974,618 haplotypes and 9,793 sites, the average query time of matches having at least one thousand sites for the single sweep long match query algorithm is 2.13 milliseconds for PBWT implementation, and 3.55 milliseconds for d-PBWT implementation (Sanaullah *et al*., 2021).

However, Naseri et al.’s and Sanaullah et al.’s query search algorithms are still not practical for genealogical search in real databases because they do not allow any mismatches when performing IBD segment detection. IBD segment may have mismatched sites due to mutation, gene conversion, or genotyping error. The exact IBD segment matching result from Naseri et al.’s or Sanaullah et al.’s algorithm may underestimate the length of the real IBD segment, which may impact the degree of relatedness inference. Thus, a new efficient haplotype query method tolerating mismatches is desired.

Here, we propose a new PBWT based IBD detection method, random projection-based identical-by-descent (IBD) detection (RaPID) query, referred as RaPID-Query. To allow mismatches while maintaining efficiency and accuracy, RaPID-Query uses the idea of multiple low-resolution PBWT panels by random projection introduced in RaPID (Naseri *et al*., 2019b). In addition, a few algorithmic innovations are introduced. First, we use x-PBWT-Query, an extended PBWT query algorithm, by simplifying Sanaullah et al.’s single sweep long match query algorithm. Second, we come up with a multi-resolution PBWT idea: while merging the results from the multiple low-resolution PBWT scans which gives approximate IBD segments, a high-resolution PBWT is run to refine the result with additional segmental reconstruction. RaPID-Query provides a promising method for fast query search. We compare the performance of RaPID-Query with the state-of-the-art all-vs-all IBD detection method Hap-IBD (Zhou *et al*., 2020a). We investigate the feasibility of the query time on large cohort dataset for RaPID-Query. We also perform the analysis of individual relatedness degree separation on IBD segments detected by RaPID-Query and compare the result with x-PBWT-Query.

## 2 Methods

The RaPID-Query method uses a new long match query algorithm, named x-PBWT-Query algorithm (i.e., extended PBWT query algorithm, by simplifying Sanaullah et al.’s single sweep long match query algorithm (Sanaullah *et al*., 2021)) to detect the IBD segment, with the random projection trait to tolerate mismatching sites in IBD segment. The resulted IBD segments from the query are refined by running long match query algorithm on the full resolution panel. The random project trait is inherited from RaPID (Naseri *et al*., 2019b). A new merging algorithm tailored to querying is proposed, which requires no additional disk space as there is no intermediate files output to the disk. Compared to PBWT query algorithms (Naseri *et al*., 2019a; Sanaullah *et al*., 2021), the IBD segments from RaPID-Query method are with high quality and allowing mismatch sites for the query search.

### 2.1 x-PBWT-Query

The x-PBWT-Query algorithm is a simplified version of Sanaullah et al.’s single sweep long match query algorithm (Sanaullah *et al*., 2021). The algorithm takes a query and a panel with *n* number of sites as the input, as well as the minimum cutoff length of the IBD segment *L* (where *L* ∈ [1, *n*]). It outputs the IBD segments found between the query individual and individual in the panel (i.e., it reports a tuple (individual_id_in_panel, IBD_start_position, IBD_end_position) for the query_individual_id). The algorithm assumes PBWT panels (i.e. prefix array *p*, divergence array *d*, and block indicator related arrays *u* and *v*) (Durbin, 2014) are pre-computed and accessible. The x-PBWT-Query algorithm pseudocode can be found in Algorithm S1, and the detail of the *getBlockIndicator*(·) function to update the match block [*f, g*) indicators *f* (or *g*) in Algorithm S1 is described in Algorithm S2, which covers both regular and boundary cases.

The Sanaullah et al.’s single sweep long match query algorithm (Sanaullah *et al*., 2021) assumes the query haplotype is virtually inserted into the panel. A virtual query indicator is used to track the query haplotype location in the panel. The algorithm adopts the set-maximal match mechanism, first introduced by Durbin (2014), to find the long match block. This is similar to the behavior of Naseri et al.’s PBWT-Query algorithm (Naseri *et al*., 2019a). If the length of the start position *e* of the set-maximal match to the current scanned site position *k* is at least the cutoff long match length *L*, a match block is formed. The newly formed match block has one haplotype only, whose location depends on the values of block indicator arrays *u* and *v*, as well as the virtual query indicator. Then, the algorithm uses the neighbor haplotypes’ divergence values to expand the match block in both directions. If such divergence value shows the length of the match between the haplotype in the match block edge and the neighbor haplotype outside of the match block edge is at least *k* + 1 − *L*, the match block is expanded. Finally, the algorithm reports matches if the matches in the match block have the opposite site value to that of its neighbor’s haplotype (i.e. a break) in the next-to-be scanned site position.

The proposed x-PBWT-Query algorithm simplifies the Sanaullah et al.’s single sweep long match query algorithm when forming the match block. Firstly, it does not use the concept of virtual insertion thus tracking the virtual query indicator is no longer needed. Secondly, it eliminates the redundant steps of evaluating the divergence values of the haplotypes if they are already in the set-maximal match block. In x-PBWT-Query algorithm, the step of updating the value of the virtual query indicator in each site is not needed. The long match block is initially formed to include all haplotypes in the set-maximal match block when the constraint “*e* = *k* + 1 − *L*” is satisfied. Figure 1 is an example of the long match block initialization process when *L* = 6. Comparing to the procedure of initializing match block in Sanaullah et al.’s single sweep long match query algorithm, where the match block starts with one haplotype, the x-PBWT-Query algorithm’s match block starts with all haplotypes in the set-maximal match block. This approach simplifies the match block build-up process by avoiding re-evaluating the divergence values of the haplotypes already in the set-maximal match block during the step of expanding the match block. Though theoretically the time complexity stays the same, it reduces the overhead in practical applications.

**Figure 1:**
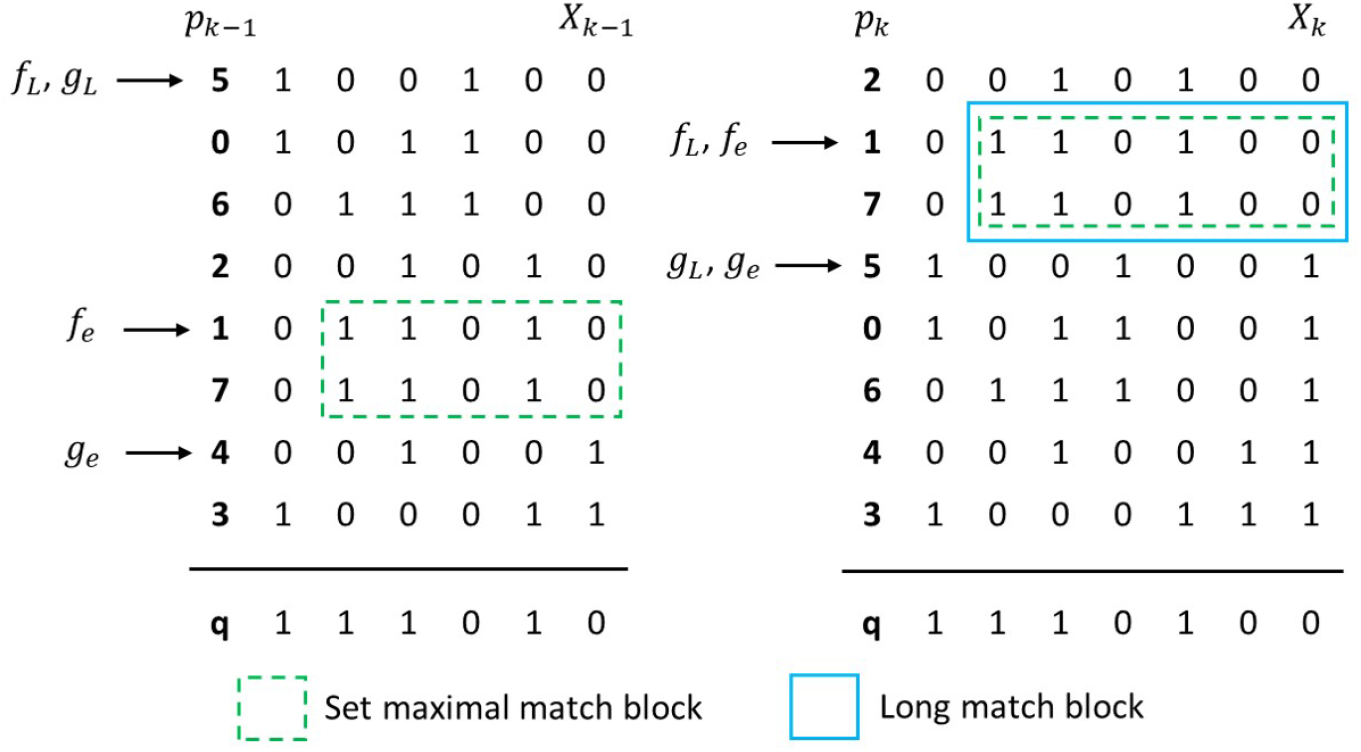
x-PBWT-Query Long Match Block Initialization Example (*L* = 6).

Additionally, The x-PBWT-Query algorithm offers a new feature: site distance tracking. This feature makes x-PBWT-Query algorithm be feasible when the cutoff long match length *L* is in either physical or genetic unit of measurement. The algorithm tracks site index *i* who is *L* away from the currently scanned site *k* while maintaining the same complexity of the algorithm. Figure 2 is an example of the updates of the site distance track index *i* in scanned site *k*. Site distance track index 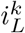 indicates the closest site having *L* centiMorgan (cM) distance away from site *k*. The site distance track index *i* is updated as the site scanning goes. Since the site scanning would never go backwards, the maximum number of the update is bounded by the number of sites (i.e. linear operation). When expanding the match block, the site distance track index *i* is marked as the start position of the matches in the match block. In Sanaullah et al.’s single sweep long match query algorithm, the start position of the matches is updated as *k* + 1 − *L*, which requires non-trivial conversion (e.g., an additional data structure holding the mapping information from genetic positions to physical positions is needed) if the given cutoff long match length *L* is in genetic distance format. The site distance track index of x-PBWT-Query algorithm facilitates the usage of the algorithm in real applications. In most real world scenarios, genetic maps are used to measure distance in chromosomes; thus, genetic length is used to represent the cutoff long match length *L*. In such case, it is very straightforward to have genetic maps directly applied to the distance track variable *i* in x-PBWT-Query algorithm, to be able to query the panel easily using the genetic distance cutoff length.

**Figure 2:**
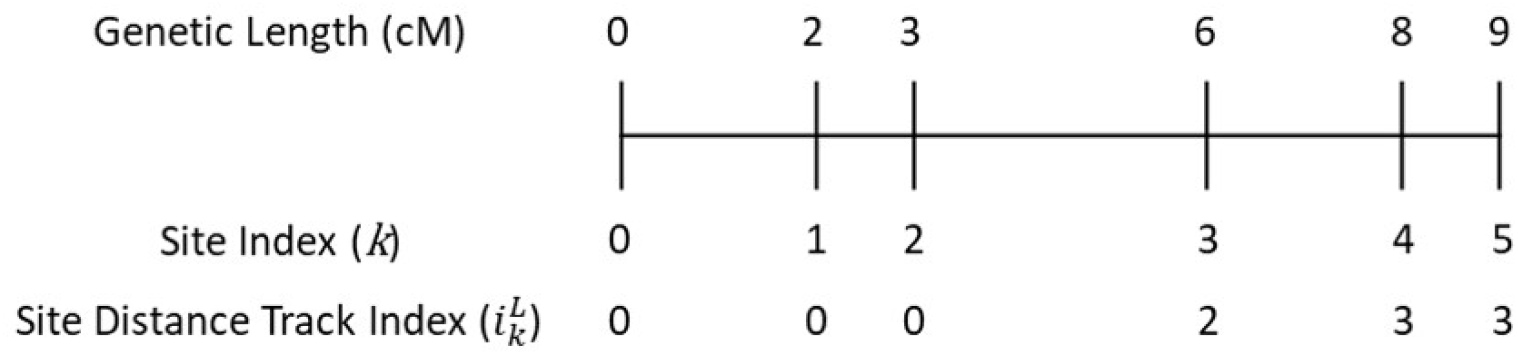
x-PBWT-Query Site Distance Track Index Example (*L* = 2).

The time complexity of the proposed x-PBWT-Query algorithm is *O*(*nm*) in worst case scenario, where *n* is the number of sites and *m* is the number of haplotypes. On average case, its time complexity is *O*(*n* + |*output*|), where |*output*| ∈ *O*(*nm*) is the number of the output IBD segments. This is equivalent to the time complexity of Durbin’s finding all set-maximal matches from a new sequence algorithm (Durbin, 2014), Naseri et al.’s PBWT-Query algorithm (Naseri *et al*., 2019a), and Sanaullah et al.’s single sweep long match query algorithm (Sanaullah *et al*., 2021), as Durbin’s procedure of finding all set-maximal matches from a new sequence, dominating the time complexity, is part of these algorithms. The empirical evaluation on the average time complexity of such procedure being *O*(*n* + |*output*|) is documented and the result is confirmed (Naseri *et al*., 2019a). The time complexity of the proposed long match query algorithm independent from the number of haplotypes *m* makes the algorithm scalable to gigantic biobank-sized or even population-scaled panels.

### 2.2 RaPID-Query

The drawback of PBWT-based long match query algorithm is not allowing mismatch sites when searching for shared IBD segments. It is common that an IBD segment contains few mismatch sites due to the event of genotyping error, gene conversion, or mutation. To have the proposed long match query algorithm allowing mismatch sites in IBD segment, the concept of random projections, originally proposed in RaPID (Naseri *et al*., 2019b), is brought into the new long match query algorithm and composes the random projection-based IBD detection (RaPID) query method, referred as RaPID-Query. Similar to RaPID, RaPID-Query divides the panel into windows and for each window a site is sampled randomly on the weight of the site frequency, to form a low resolution panel. RaPID-Query generates multiple low resolution panels and then, the x-PBWT-Query algorithm is run on each low resolution panel to find exact matched segments. The next step is to combine result segments from all the runs. A new IBD segment merging method tailored to querying is proposed. Different from RaPID, the proposed merging method is scalable to large number of detected IBD segments as there is no intermediate files output to the disk, eliminating the situation that poor disk I/O performance hampers the run time. Finally, the x-PBWT-Query algorithm is run on the original resolution panel to get the IBD segments in full resolution. The full resolution IBD segments are used to refine the boundaries of low resolution IBD segments, with the cases of stitching IBDs if their distance in chromosome region is within the allowed gap *g*_*max*_. Figure 3 shows the entire RaPID-Query workflow. The left section shows the pre-process part of the RaPID-Query method. The population panel and associated PBWT panels and their sub panels are calculated in advance. The right section shows the query search part of the RaPID-Query method. The associated sub queries are generated based on the provided individual query and search is performed quickly thanks to all the pre-processed panels and sub panels.

**Figure 3:**
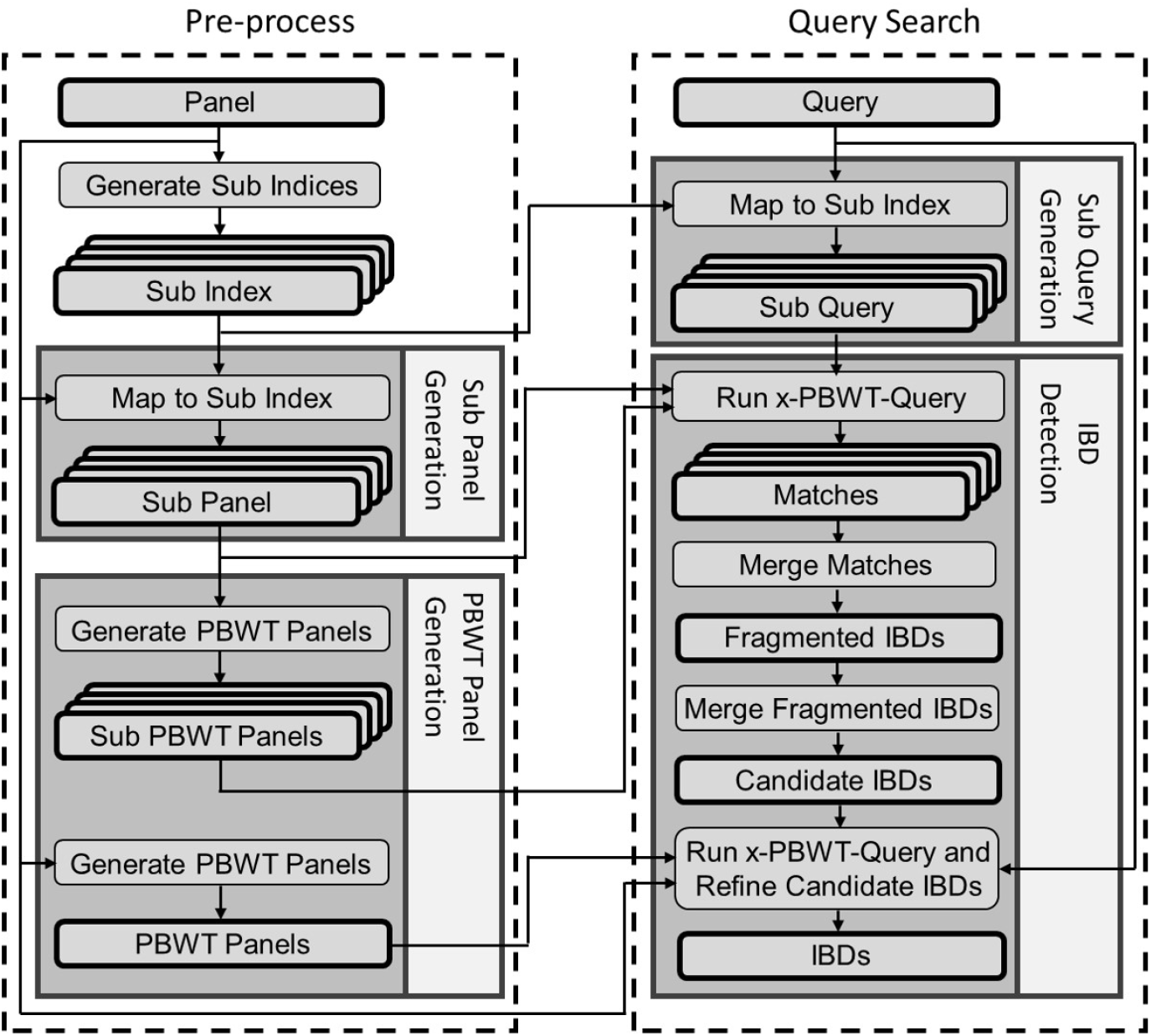
RaPID-Query Workflow.

#### 2.2.1 Query with Random Projection

The first part of the method is to produce matches for the query allowing mismatch sites. Assuming long matches with minimum match cutoff length *L* are to be found on a panel with total number of *n* sites and *m* haplotypes for a query haplotype with the same *n* sites. During the pre-process stage, the PBWT panels (i.e., prefix array *p*, divergence array *d*, and block indicator related arrays *u* and *v*) as well as the minor allele frequencies of each site are calculated based on the input site panel. Then, the random projection approach is used to create down-sampled site panels from the original site panel. The site panel is equally divided into 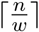 sections, where each section’s window size is *w*. The sub index is sampled from the index of the site panel, one site index per section, by using weighted random sampling. The probability of each site index getting selected is determined by the minor allele frequency of the site in the section. The larger the minor allele frequency the site has, the more chance of such site is getting selected. The sub index is formed with 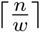 site indices. For each random projection run, one sub index is needed. Thus, there are *r* sub indices generated in the pre-process stage where *r* is the number of random projection runs. Each sub index is correlated to a sub panel. Next, the sub PBWT panels are created for each sub panel and ready for the queries. In the query search stage, *r* sub queries are generated by mapping the input query site panel to the sub indices. Then, x-PBWT-Query algorithm is applied on each sub query and correlated sub panel and sub PBWT panels, to output the detected matches. The detected matches are rescaled from the resolution window size to the original panel and query size. Specifically, a low resolution match (*id*, [*h*′, *t*′]) found from the low resolution panel with the window size *w*, is rescaled to the original resolution as (*id*, [*wh*′, *w*(*t*′ + 1) −1]) (or (*id*, [*h, t*])), where *id* is the haplotype identification number who has the match with the query haplotype, *h*′ (or *h* = *wh*′) is the head or start position of the match and *t*′ (or *t* = *w*(*t*′ + 1) − 1) is the tail or end position of the match, inclusively. Figure S1 shows an example of query with random projection runs.

#### 2.2.2 IBD Segment Identification

The second part of the method is to merge and refine the detected matches found from x-PBWT-Query results. To identify IBDs from the detected matches, firstly, the detected matches are grouped by the end position of the match. Secondly, the fragmented IBDs are identified as the match having the *c*-th smallest start position in each group, and the other matches are discarded. Thirdly, the candidate IBDs are formed by merging the fragmented IBDs, if two fragmented IBDs have overlap. The candidate IBD is formed as the union of the two fragmented IBDs. Lastly, the candidate IBDs are refined by comparing to the exact matched IBDs from the original resolution run using x-PBWT-Query algorithm. The boundaries of candidate IBD who has overlap with the exact matched IBD is extended or trimmed according to the overlapped exact-matched IBD. If the distance between two candidate IBDs is within the maximum gap threshold *g*_*max*_, the IBD is formed from stitching the two candidate IBDs. This is the special case of IBD boundary correction. Figure 4 is an IBD identification example of a haplotype pair with *c* = 2 and *g*_*max*_ = 2*cM*. It shows there are six groups of detected matches (i.e., G1-G6) after grouping the end position of the match. Then, five matches having the second smallest start position in each group, are identified as the fragmented IBDs (i.e., G1-G2, G4-G6). The match in G3 is discarded as there is no match having the second smallest start position existing in the group. Then, the fragmented IBDs are merged if they have overlap across the groups. In the example, three candidate IBDs are formed: one from group G1, one from group G2, and the other one from the union of three fragmented IBDs from group G4-G6 since they have overlaps. Finally, the three candidate IBDs are refined into two IBD segments: the candidate IBDs from group G1 and G2 are stitched together, as the gap between them in chromosome region is less than 2 cM; the candidate IBD from the group G6 is trimmed on both ends, to be aligned with the boundaries of the overlapped exact matched IBD segment.

**Figure 4:**
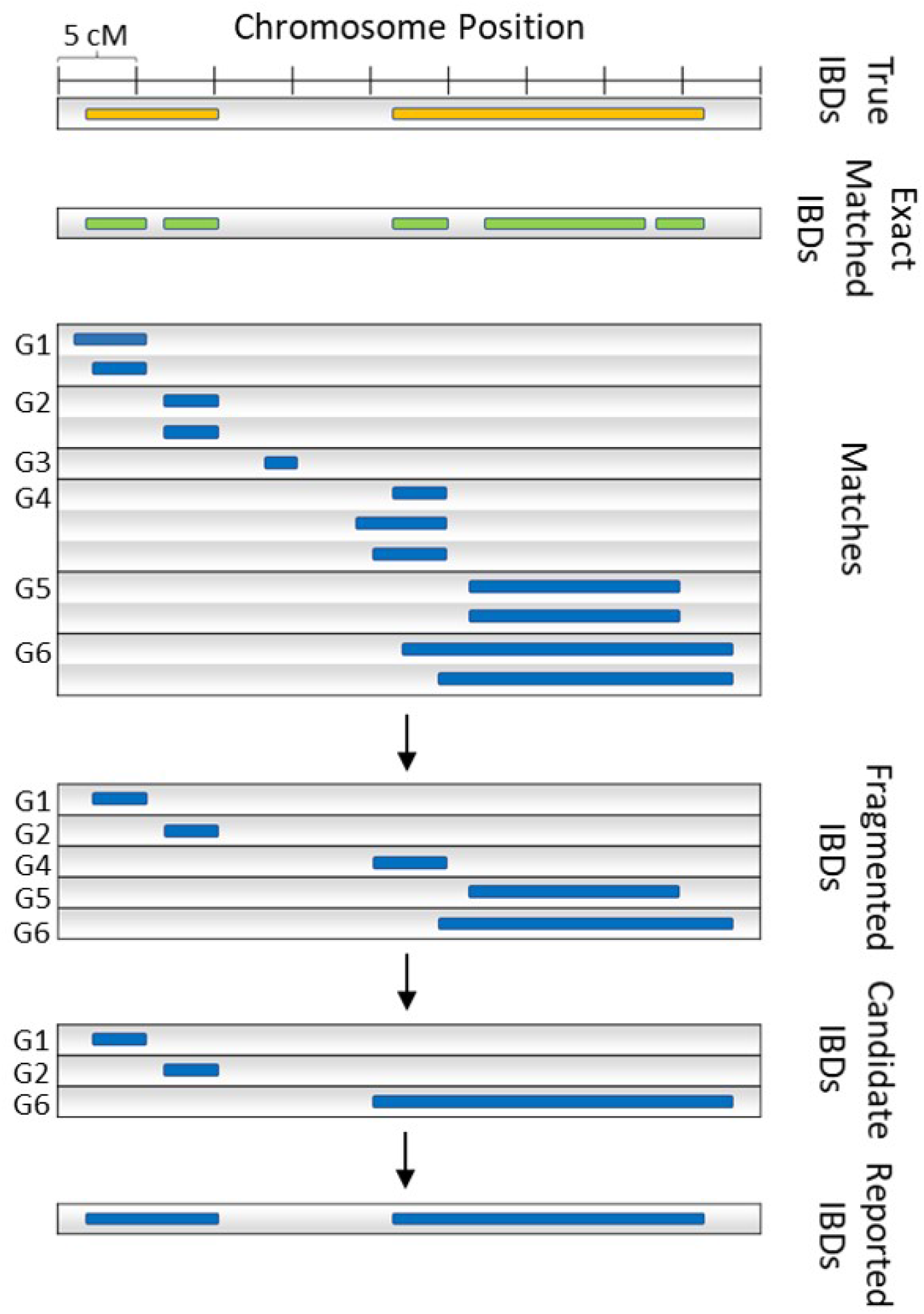
Merge and Refine Example of a Haplotype Pair (*c* = 2, *g*_*max*_ = 2*cM*). Groups containing no fragmented or candidate IBD segments are omitted.

To group the end positions of the matches from the random projection runs, each detected matches (*id*, [*h, t*]) is efficiently stored in an array of hash tables. In particular, a PBWT bucket data structure *B*_*a*_ is introduced to store and merge detected matches from the queries among random projected panels. *B*_*a*_ is an array of size *n*, whose index is *t*, indicating the end position of the match. It stores information of all matches found ending at position *t* inclusively, which is a hash table data structure whose key is the matched haplotype *id* and value is a list *H* storing the start position *h* and *count*, indicating the number of identical matches found, for all candidate matches found having the same haplotype *id* and ending at the same position *t*.

The way to collect all matches found from PBWT queries results is to collect matches per site, starting from site 0 to site *n* − 1 from all random projection runs in the same resolution group. The matches found from all runs at site *t* are collected and stored in PBWT bucket *B*_*a*_, before each PBWT query moves to next site. See Algorithm S3. For each newly found match (*id*_*new*_, [*h*_*new*_, *t*]) from the runs, first it is merged with existing matches ending at the same current site *t* in bucket *B*_*a*_: the new match is inserted in *B*_*a*_ if it is not found; otherwise, for existing matches, if the match interval of the new match encompasses the match interval of an existing match in *B*_*a*_ for the same sample, the existing match’s *count* value is incremented by one. Second, a search needs to be performed to merge the new match with existing matches ending at previous sites (i.e., site value is less than *t*). The search range is from the starting position of the longest match found at site *t* to its ending position *t* (i.e., the coverage is [*h*_*min at t*_, *t*)). Since all matches are rescaled and stored in original size bucket *B*_*a*_ (which contains *n* site slots), they can be found only in every *w* site slots. When searching from site *t* back to site *h*_*min at t*_, instead of checking every site, each time the search can jump window size length *w* distance. This can avoid unnecessary searches. The search starts from site *t* − *w* to search the matches met the criteria, and the next group of matches to be searched is located on *w* distance away from the current one. The search criteria is similar as the one searching matches ending at *t*: if the interval of the new match encompasses the interval of the existing matches, their *count* values are incremented.

The complexity of collecting matches from runs on randomly projected panels in the same resolution level is bounded by the total number of matches from all runs and the number of search times to find and merge match with existing matches. For the part of merging matches ending at site *t*, there are constant operations to fetch matches from the list storing match starting positions *h* in *H*, since in most cases there is only a small number of matches found per site. The maximum size of the list is *r*, which is a constant number, as for each run it is not possible to found a match ending at the same position twice. If *C*_*low*_ is a set containing all matches found from the random projected low resolution panels, the time complexity of the part of merging matches ending at site *t* is the number of matches found from all runs, i.e. *O*(|*C*_*low*_|). For the part of merging matches ending at site less than *t*, the complexity depends on the number of search times to find and merge match with existing matches, i.e., the sum of distances that the search performed backwards at each site. For each site *t*, such number is 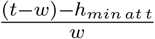. Thus, for all sites, this number is 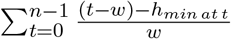. Let *l* = *t* − *h*_*min at t*_ denote the length of the longest match found at site *t*. The number of search times can be written as 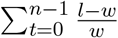, which indicates the time complexity of collecting matches from runs in random projected low resolution panels is 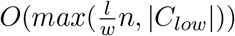. The space complexity of collecting matches from runs in same resolution panels is *O*(*n* + |*C*_*low*_|), as the PBWT bucket *B*_*a*_ is an array of size *n* containing matched haplotypes *id* and starting positions *h*, whose space is bounded by the total number of matches found from all runs, |*C*_*low*_|.

The fragmented IBDs are identified from the detected matches by filtering out the false positive matches. This step only requires a simple traversal on PBWT bucket *B*_*a*_ which stores all detected matches. Algorithm S4 shows the detail of the filtering. The algorithm traverses *B*_*a*_ while checking the *count* value of each match. If the *count* value is less than the number of count success value *c*, the match is considered as a false positive and removed from *B*_*a*_. After the traversal, all false positive matches are filtered out and the remaining matches in *B*_*a*_ are potential matches. The time complexity of filtering out false positive matches is *O*(|*C*_*low*_|), as this algorithm traverses each match exactly one time. The space complexity is *O*(*n* + |*C*_*low*_|).

The candidate IBDs are formed by merging overlapped fragmented IBDs in *B*_*a*_. The overlapped fragmented IBDs are considered as two fragmented IBDs: (*id*, [*h*_1_, *t*_1_]) and (*id*, [*h*_2_, *t*_2_]), if and only if *t*_1_ ≥ *h*_2_ or *t*_2_ ≥ *h*_1_. Algorithm S5 checks fragmented IBDs in the order of their ending positions (i.e. from site 0 to *n* − 1) and merges or appends them if any overlapped segments are found. For each fragmented IBD, the algorithm checks all other fragmented IBDs ending at the same site *t* as well as those ending at site less than *t*, to find if they meet the criteria to be merged or appended. Firstly, it checks fragmented IBDs ending at site *t*: for each fragmented IBDs stored in *H*, only match (*id*, [*h*_*min at t*_, *t*]) is kept since it encompasses all other fragmented IBDs ending at site *t*. All other fragmented IBDs are removed from *H*. Secondly, it checks fragmented IBDs ending at site less than *t*: it visits sites backwards from *t* − *w* site to *h*_*min at t*_ − 1 for every *w* sites. Site *h*_*min at t*_ − 1 is a boundary case (i.e., the fragmented IBD’s ending position is adjacent to the current fragmented IBD’s starting position). If such fragmented IBD is found, the current fragmented IBD is appended to the found fragmented IBD as one. If a fragmented IBD (*id*, [*h*_*prev*_, *t*_*prev*_]) is found within the range [*h*_*min at t*_, *t*], the fragmented IBD (*id*, [*h*_*min at t*_, *t*])’s starting position is update as *min*(*h*_*min at t*_, *h*_*prev*_). The fragmented IBD (*id*, [*h*_*prev*_, *t*_*prev*_]) is removed from *H*.

The complexity of merging overlapped fragmented IBDs is bounded by the total number of fragmented IBDs in *B*_*a*_ and the number of search times to find and merge overlapped fragmented IBDs. For the part of merging fragmented IBDs ending at site *t*, keeping fragmented IBD (*id*, [*h*_*min at t*_, *t*]) and removing all other fragmented IBDs costs 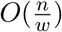. For the part of merging fragmented IBDs ending at site less than *t*, the traversal from *t* − *w* site to *h*_*min at t*_ − 1 costs 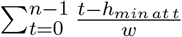 number of search times. Using the same notation as previously described: *l* = *t − h*_*min at t*_, which is the length of the longest fragmented IBD found at site *t*, the time complexity of the step is 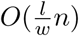. Though *h*_*min at t*_ may get updated as *h*_*prev*_, if a fragmented IBD (*id*, [*h*_*prev*_, *t*_*prev*_]) is found at site *t*_*prev*_ and *h*_*prev*_ *< h*_*min at t*_, the search stops after the update. Since site *t*_*prev*_ is processed before site *t*, either no fragmented IBD, or just one fragmented IBD (*id*, [*h*_*prev*_, *t*_*prev*_]) exist. If it exists, the starting position of the fragmented IBD *h*_*prev*_ is updated previously as well and no other *h* value smaller than *h*_*prev*_ exists prior to site *t*_*prev*_ that such fragmented IBD has any overlaps with the fragmented IBD (*id*, [*h*_*min at t*_, *t*]). Thus, the time complexity is 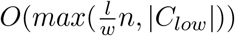. Same as previous, the space complexity of the algorithm is *O*(*n* + |*C*_*low*_|).

The IBDs are called from the refinement of the candidate IBDs with the guidance of the exact matched IBDs from the run on full resolution panel. A refine-on-the-fly approach is used: while running the long match query on full resolution panel, if an exact matched IBD is found, and if it has overlap with a candidate IBD, it is used to refine such candidate IBD; otherwise it is discarded. Algorithm S7 shows the detail of the refinement algorithm. The start and end boundary of the calling IBD lines up with the exact matched IBD found from the run on full resolution panel, who has overlapped segment with the candidate IBD found from runs on low resolution panels. Additionally, the algorithm is allowed to have a gap *g*_*max*_ in a calling IBD. Similar to the maximum gap length idea that Hap-IBD uses (Zhou *et al*., 2020a), such gap is considered as genotyping error, mutation, or gene conversion. This is to rectify the small gap on the IBD by stitching two candidate IBDs whose distance is within the gap distance *g*_*max*_ on a chromosome region, when such gap is not picked up by random projection runs. The length of the IBD to be reported needs to be at least *L* after the refinement.

To efficiently refine the candidate IBDs, a PBWT bucket data structure *B*_*m*_ is used to store and refine the candidate IBDs originally found from random projection runs in low resolution panels. *B*_*m*_ is a hash table data structure whose key is the matched haplotype *id* and value is a stack *S* storing the location of the candidate IBD found from runs in low resolution panels (i.e. (*h, t*)) in an ascending order of the start position *h*, and the location of the IBD segment to be reported, resulted from the last exact matched IBD found from the run in full resolution panel (i.e. *R* = (*h*_*R*_, *t*_*R*_)). Algorithm S6 shows the conversion of the candidate IBD form from *B*_*a*_ to *B*_*m*_ with a sorted order of the candidate IBD locations in a linear fashion.

If *C*_*low*_ is a set containing all candidate IBDs found in random projected low resolution panels and *C*_*full*_ is a set containing all exact matched IBDs found in full or original resolution panel, the time complexity of the refinement algorithm is 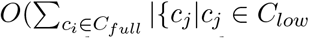, *and c*_*i*_.*id* == *c*_*j*_.*id*}|); however, the candidate IBDs found from runs in low resolution panels are sorted by their locations in *B*_*m*_, and the exact matched IBDs found from the run in full resolution panel appear following the order of the site index, the time complexity is not a Cartesian product of the two sets. Instead, the actual time complexity is *O*(*max*(*n*, |*C*_*low*_|+|*C*_*full*_|)) as each match is evaluated only once. The space complexity is *O*(*n* + |*C*_*low*_|) as the exact matched IBDs found from the run in full resolution are not stored during the refinement.

## 3 Results

To test out the performance of RaPID-Query method, sets of experiments are performed on simulated data and real-world data. The experiments on the simulated dataset show the feasibility of IBD segment detection using RaPID-Query method. The experiments on the real-world dataset prove the practicability of using RaPID-Query method to perform IBD segment based genealogical searches. The machine used for the experiments has Intel Xeon Gold 5215 2.50 GHz processor with 3 terabytes of RAM.

### 3.1 Dataset

A 2,000-haplotype sequencing dataset was simulated to test the performance of RaPID-Query method. The simulation used the out-of-Africa demographic model (Gutenkunst *et al*., 2009) via msprime v1.0.1 (Kelleher *et al*., 2016) with the parameters provided by stdpopsim library (Adrion *et al*., 2020). The chromosome 20 genetic map in GRCh38 coordinates from deCODE genetics (Hall-dorsson *et al*., 2019) was used as the recombination map. The mutation rate was set to 1.38e-08 as it was the same constant rate in simulated dataset tested by the state-of-the-art all-vs-all IBD detection method: Hap-IBD (Zhou *et al*., 2020a). This mutation rate also falls within the most recent estimated mutation rate 95% confidence interval: [1.02e-8, 1.56e-8] (Tian *et al*., 2019). The simulated dataset consists 1,000 individuals, sampled from the European population. The sites with multi-allelic values or minor allele frequency less than or equal to 1% were filtered out from the original dataset, yielding total 92,296 bi-allelic sites. An 0.04% genotyping error was added to the dataset as it is the average error rate in TOPMed sequencing dataset (Taliun *et al*., 2021). The ground truth of IBD segment was identified as the contiguous segment where the two individuals have the same most recent common ancestor (MRCA) in the simulated coalescent trees. By adopting the same efficient process as Browning et al. did (2013), the trees were sampled per 5,000 genome position and the minimum genetic length of a true IBD segment calling is 1 cM.

To show the detected IBD segments from RaPID-Query are robust for genealogical search, the UK biobank SNP-array genotyping dataset (Bycroft *et al*., 2018) was used as the real-world testbed. All autosomal chromosomes containing 487,409 individuals with 658,719 sites were used to show that the IBD segments have the power of separating different degrees of relationships. The UK biobank dataset contains the ground truth of the relatedness of individuals up to third degree, measured as part of the UK biobank study and presented in the form of the kinship coefficient. The estimation of such kinship coefficient is generated by the kinship coefficient, Kinship-based INference for Genome-wide association studies (KING) software (Manichaikul *et al*., 2010). The actual degree of relatedness is determined by the ranges where the kinship coefficient resides in: first degree: (0.177, 0.354], second degree: (0.0884, 0.177], and third degree: (0.0442, 0.0884] (Manichaikul *et al*., 2010).

To demonstrate the robustness of distinguishing the different degrees of relatedness of individuals up to fourth degree using the detected IBD segments from RaPID-Query, 22 chromosomes were simulated. The simulated dataset was created by an in-house simulation program mimicing the UK biobank SNP-array genotyping dataset. 1,000 unrelated individuals from the UK Biobank were randomly selected and the population size at each generation was set to 1,000. The genetic data of the last four generations comprising 4,000 individuals including their relationships were extracted. To make the simulated dataset more realistic, an 0.13% genotyping error was added to the simulated dataset as it is the mean discordance error rate in UK biobank SNP-array genotyping data (Bycroft *et al*., 2018).

### 3.2 Power and Accuracy

The power and accuracy of RaPID-Query is demonstrated by two measurements: false negative rate and false positive rate. In this set of experiments, RaPID-Query with different sets of refinement parameters were examined against the x-PBWT-Query method. The refinement parameter, indicating the cutoff genetic length of exact match IBDs run in original resolution panel, was used as the suffix of the RaPID-Query method name (e.g., RaPID-Query-0.5 indicates the match cutoff length in original resolution panel is 0.5 cM). The other parameters were inherited from RaPID (Naseri *et al*., 2019b) and the values were set on an empirical basis. The results of all 2,000 haplotype queries were merged and averaged, in order to compare the results to the non-query-based all-vs-all state-of-the-art method Hap-IBD (Zhou *et al*., 2020a). The parameters of each methods used for the performance analysis are in Table 1.

**Table 1:** Parameters used for performance analysis of simulated dataset with minimum 2 cM IBD segment length and minimum 200 markers

| Method | Parameters |
| --- | --- |
| x-PBWT-Query | -d 2.0 -lm 100 |
| RaPID-Query-0.5 | -w 13 -r 5 -c 1 -d 2.0 -lm 200 -dh 0.5 -lmh 100 -dg 2.0 |
| RaPID-Query-1.0 | -w 13 -r 5 -c 1 -d 2.0 -lm 200 -dh 1.0 -lmh 100 -dg 2.0 |
| RaPID-Query-2.0 | -w 13 -r 5 -c 1 -d 2.0 -lm 200 -dh 2.0 -lmh 100 -dg 2.0 |
| Hap-IBD (v1.0) | min-seed=2.0 min-extend=1.0 min-output=2.0 max-gap=1000 min-markers=100 min-mac=2 |

#### 3.2.1 False Negative Rate

The measurement of false negative rate is defined as the average proportion of true IBD segments not having been covered by reported IBD segments, i.e., the sum of proportions of true IBD segments not overlapped with reported IBD segments over the number of true IBD segments:

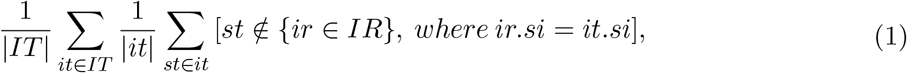

where *IT* is true IBD set, *it* is true IBD, *st* is site of true IBD, *IR* is reported IBD set, *ir* is reported IBD, and *si* is the sampled individual ID.

Figure 5 shows the false negative rates of four 1-vs-all IBD detection methods: x-PBWT-Query, RaPID-Query-0.5, RaPID-Query-1.0, RaPID-Query-2.0, and one all-vs-all IBD detection method: Hap-IBD. Overall, as the genetic length of true IBD increases, the false negative rate decreases. The x-PBWT-Query method has the worst false negative rate as it does not allow mismatches. It misses some of the IBD segments which are not exact matches and caused by genotyping errors, mutations, or gene conversions. All other methods have better false negative rates as they allow mismatches during the IBD calling. For RaPID-Query methods, the smaller the refinement parameter value is, the better the false negative rate the method has. The refinement parameters 0.5, 1.0, and 2.0 in figure are the cutoff genetic length of exact match IBDs run in original resolution panel. In this example, RaPID-Query-0.5 has the best false negative rate as it detects the largest number of exact match IBDs, which helps extending and stitching the candidate IBDs. The false negative rate of Hap-IBD is between that of RaPID-Query-1.0 and RaPID-Query-2.0 for all the cases, indicating the false negative rate of RaPID-Query is competitive to the state-of-the-art all-vs-all IBD detection method. If the cutoff genetic length of IBD is 7 cM or above, all RaPID-Query methods showing in Figure 5 has very small false negative rate (i.e., less than 10%). This shows RaPID-Query is outstanding from the power of IBD segment detection perspective when performing genealogical searches.

**Figure 5:**
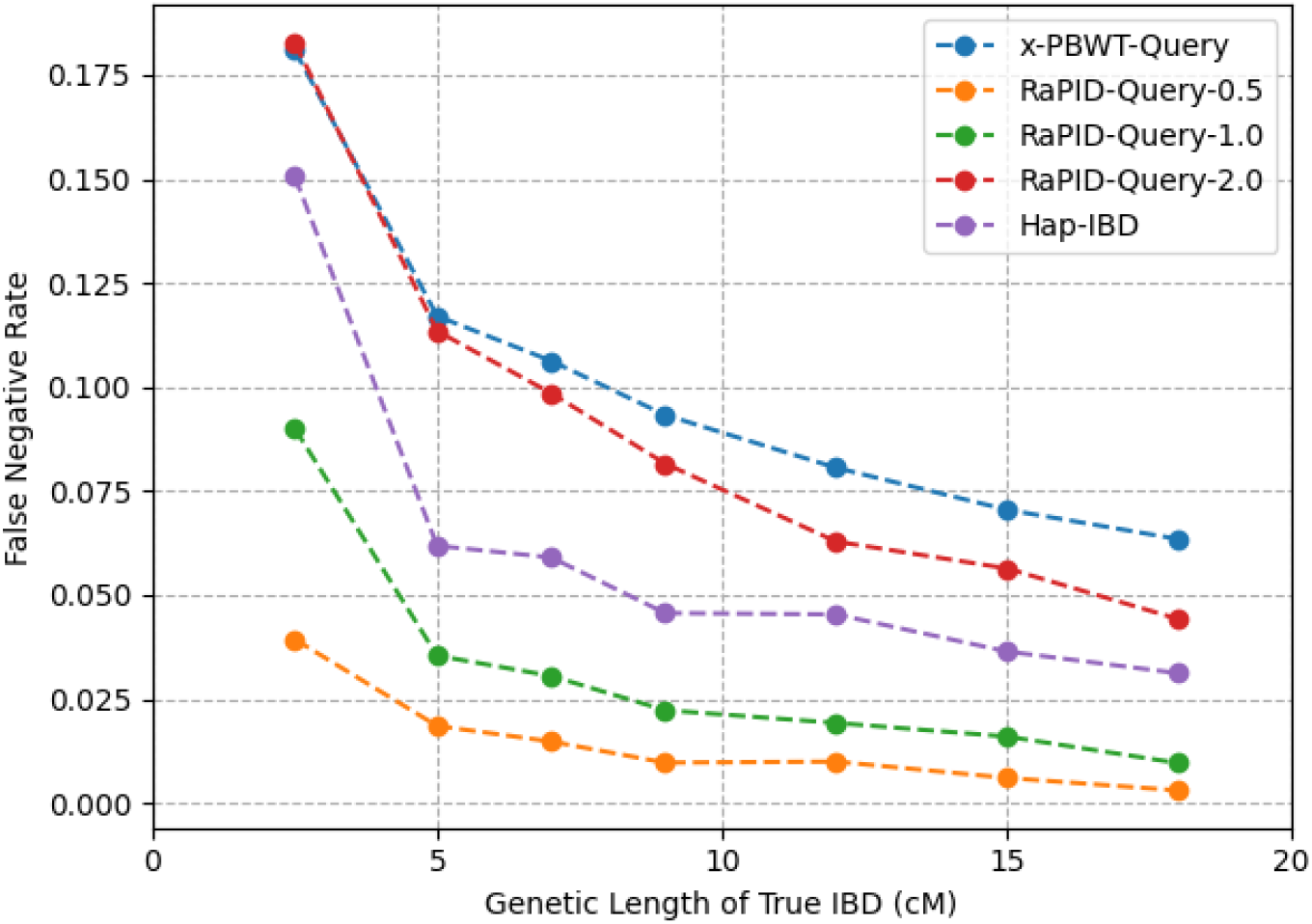
False Negative Rates. True IBD segments with length ≥ 2.5 cM were assigned into bins of 2.5–5, 5–7, 7–9, 9–12, 12–15, 15-18, and ≥ 18 cM according to their genetic length. The false negative rate is the proportion of true IBD segments in a bin that are not covered by any reported IBD segment ≥ 2 cM.

#### 3.2.2 False Positive Rate

The measurement of false positive rate is defined as the average proportion of reported IBD segments not being covered by true IBD segments, i.e., the sum of proportions of reported IBD segments not overlapped with true IBD segments over the number of reported IBD segments:

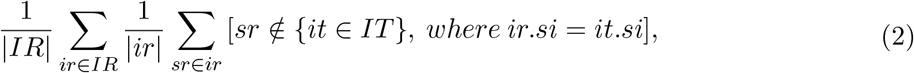

where *IR* is reported IBD set, *ir* is reported IBD, *sr* is site of reported IBD, *IT* is true IBD set, *it* is true IBD, and *si* is the sampled individual ID.

Figure 6 show the false positive rates of four 1-vs-all IBD detection methods: x-PBWT-Query, RaPID-Query-0.5, RaPID-Query-1.0, RaPID-Query-2.0, and one all-vs-all IBD detection method: Hap-IBD. Overall, as the genetic length of reported IBD increases, the false positive rate decreases. Though the false positive rates of all five methods are close in almost all cases, there is difference among the methods. The x-PBWT-Query method has the best false positive rate as it does not allow mismatches. Other methods may detect inexact match IBD segments which could be false positive ones. For RaPID-Query methods, the larger the refinement parameter value is, the better the false positive rate the method has. The refinement parameters 0.5, 1.0, and 2.0 in figure are the cutoff genetic length of exact match IBDs run in original resolution panel. In this example, RaPID-Query-2.0 has the best false positive rate as it detects the smallest number of exact match IBDs, which prevents over-extending the candidate IBDs resulted from the random projection runs. The false positive rate of Hap-IBD is similar to that of RaPID-Query-0.5 except it is slightly better than RaPID-Query-0.5 method on 2-5 and 15-18 cM ranges of the cutoff genetic length of reported IBDs. This indicates that the false positive rate of RaPID-Query is competitive to the state-of-the-art all-vs-all IBD detection method. If the cutoff genetic length of IBD is 7 cM or above, all RaPID-Query methods showing in Figure 6 has very small false positive rate (i.e., less than 2%). This shows RaPID-Query is outstanding from the accuracy of IBD segment detection perspective when performing genealogical searches.

**Figure 6:**
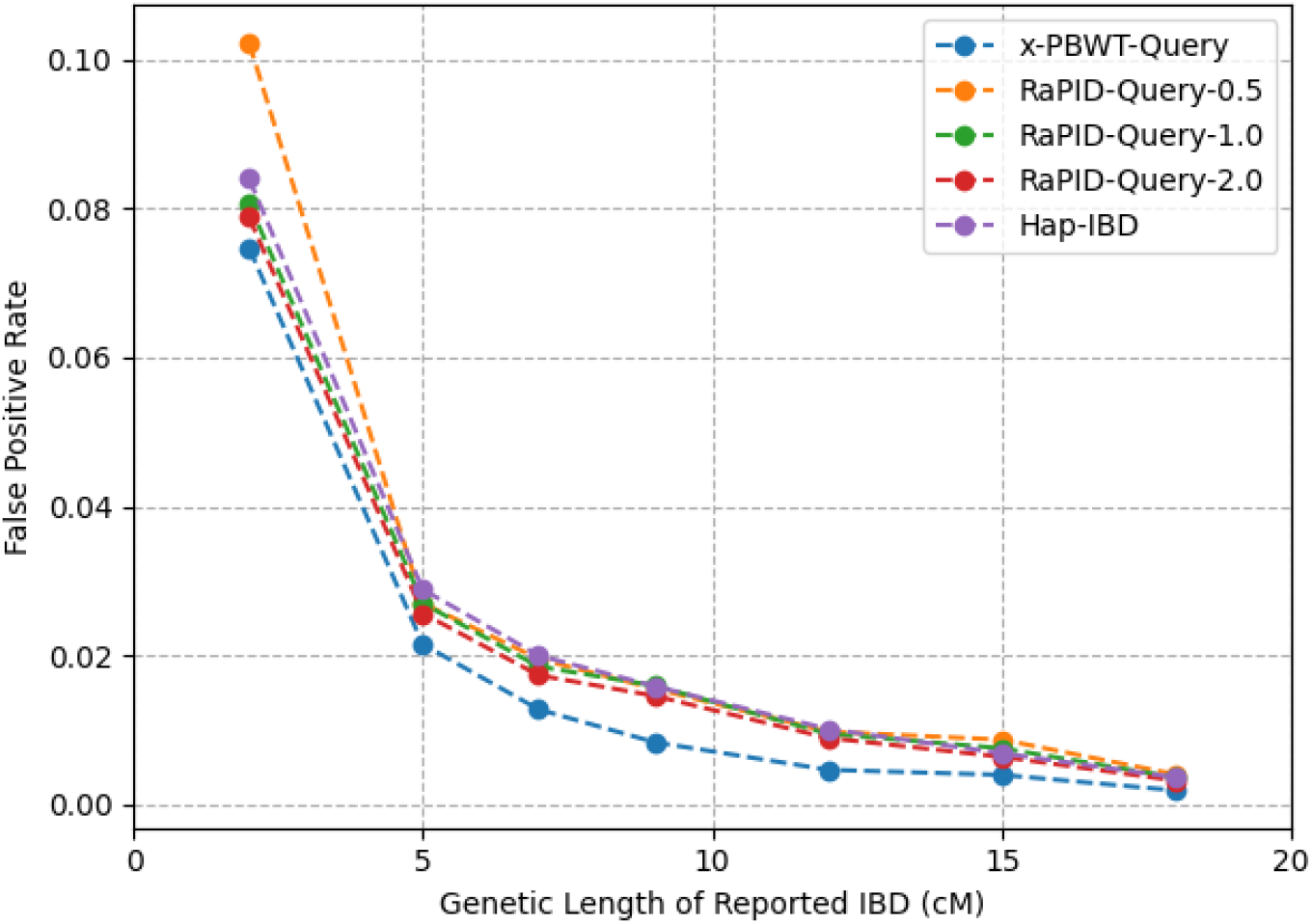
False Positive Rates. Reported IBD segments with length ≥ 2 cM were assigned into bins of 2–5, 5–7, 7–9, 9–12, 12–15, 15-18, and ≥ 18 cM according to their genetic length. The false positive rate is the proportion of reported IBD segments in a bin that are not covered by any true IBD segment ≥ 1.5 cM.

### 3.3 Run Time

The run time of RaPID-Query was tested with different sets of refinement parameters (same used in correctness experiment) with x-PBWT-Query on the UK biobank SNP-array genotyping chromosome 20 dataset, containing 974,818 haplotypes with 17,197 sites (Bycroft *et al*., 2018). The GRCh37 human genome assembly from HapMap Phase II project (Consortium. and centres: Perlegen Sciences., 2007) were used to determine the length of IBDs detected from each autosomal chromosome, as the positions of the UK biobank SNP-array genotyping calls are in GRCh37 alignments. For each query method, 400 haplotype queries were run with 7 cM and 700 sites as the cutoff IBD segment length.

Table 2 shows the average CPU time of one query against a large panel for RaPID-Query-0.5, RaPID-Query-1.0, RaPID-Query-2.0, and x-PBWT-Query method. x-PBWT-Query is the fastest method among all, as it is not involved in any multiple runs and merges as RaPID-Query method does. For RaPID-Query method, the value of the refinement parameter is inversely proportional to the average query time. This is expected as RaPID-Query needs more time to refine the detected IBD segments as the required refinement precision increases. Even so, the RaPID-Query-0.5 method whose refinement precision is up to 0.5 cM is fast: it only takes 839.63 milliseconds on average to complete a query and write the resulted IBD segments to a file.

**Table 2:** Computation times of 400 queries against UK biobank chromosome 20 dataset with minimum 7 cM IBD segment length and minimum 700 markers

| Method | Average CPU Time (millisecond) | Standard Deviation | Average Wall Time (millisecond) | Standard Deviation |
| --- | --- | --- | --- | --- |
| RaPID-Query-0.5 | 839.63 | 166.26 | 864.47 | 153.69 |
| RaPID-Query-1.0 | 211.73 | 64.77 | 219.46 | 57.09 |
| RaPID-Query-2.0 | 39.58 | 20.06 | 48.61 | 24.05 |
| x-PBWT-Query | 11.80 | 8.02 | 18.32 | 10.10 |

### 3.4 RaPID-Query for Genealogical Analysis

To show the detected IBD segments from RaPID-Query are robust for genealogical search, a similar analysis of sum-of-IBD-length-based relatedness degree separation (Naseri *et al*., 2019a) was conducted on UK biobank and simulated datasets. For UK biobank dataset, the queries contain 200 randomly-selected individuals having at least third degree of relatedness with each other, and 1,000 randomly-selected unrelated individuals (excluding those 200 individuals having at least third degree of relatedness with each other). For the simulated dataset, the queries are 200-randomly-selected individuals. The genealogical search minimum cutoff length of an IBD calling for both datasets is 7 cM and 700 markers, as it is the conventional minimum match threshold values currently used by major direct-to-consumer (DTC) companies such as 23andMe (https://customercare.23andme.com/hc/en-us/articles/212170958-DNA-Relatives-Detecting-Relatives-and-Predicting-Relationships). RaPID-Query-2.0 was used since from previous experiments, it was the fastest method with comparable false negative and false positive rates among the methods.

Figure 7 shows the probability distributions of total length of IBDs between individual pairs on UK biobank dataset. The probability distributions of four degrees (first-degree, second-degree, third-degree, and unrelated) in the results from x-PBWT-Query method (Figure 7A) and from RaPID-Query-2.0 method (Figure 7B) are distinguishable. The probability distributions resulted from x-PBWT-Query method have smaller means and larger standard deviations than those resulted from RaPID-Query-2.0 method (See Table S1). This tells that RaPID-Query-2.0 identifies more robust IBDs than that of x-PBWT-Query, as RaPID-Query-2.0 allows mismatched sites, which makes the reported IBDs more close to the true IBDs. On the other hand, with no support on tolerating mismatched sites, x-PBWT-Query misses many IBD segments which flattens the probability distributions of total length of IBDs between individual pairs and makes them all shifted towards right in x axis.

**Figure 7:**
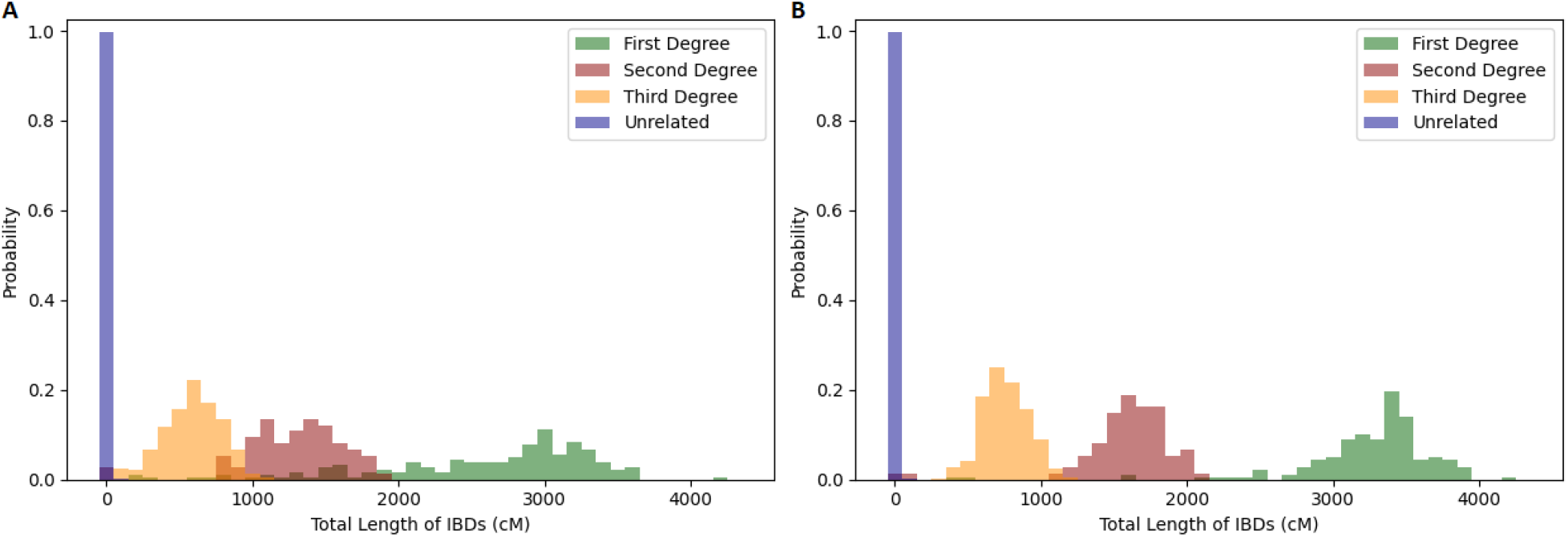
Probability Distributions of Sum of Length of IBDs on UK Biobank Dataset. (A) x-PBWT-Query. (B) RaPID-Query-2.0.

To quantify the relatedness degree separation result, the area under the receiver operating characteristic (ROC) curve (AUC) values between each distribution pairs were calculated. The AUC values of sum of length of IBDs on UK biobank dataset are shown in Table 3 (The ROC curves are in Figure S2). RaPID-Query-2.0’s AUC value between first- and second-degree relationships is 98.27%, and the AUC values between second- and third-degree relationships and third-degree and unrelated relationships are 97.28% and 100.00%, respectively. This indicates that IBD segments identified by RaPID-Query-2.0 is capable of being used to infer up-to-third-degree familial relatedness between any individual pair in large real-world biobank. Additionally, it is observed that all AUC values calculated from the sum of length of IBDs using RaPID-Query-2.0 are better than those using x-PBWT-Query, which means RaPID-Query-2.0 method performs better relatedness degree separation than x-PBWT-Query method does.

**Table 3:**
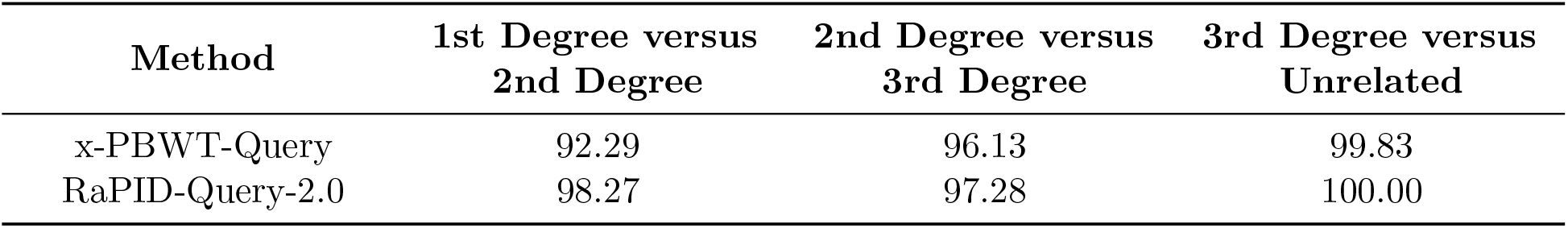
x-PBWT-Query versus RaPID-Query-2.0: Area Under Curve (%) of Sum of Length of IBDs on UK Biobank Dataset

The high-quality IBD segments from RaPID-Query-2.0 is able to separate the fourth-degree distribution from others well in simulated dataset. Figure 8 shows the probability distributions of total length of IBDs between individual pairs on simulated dataset. The probability distributions of five degrees (first-degree, second-degree, third-degree, fourth-degree, and unrelated) in the results from x-PBWT-Query method (Figure 8A) and from RaPID-Query-2.0 method (Figure 8B) are distinguishable.

**Figure 8:**
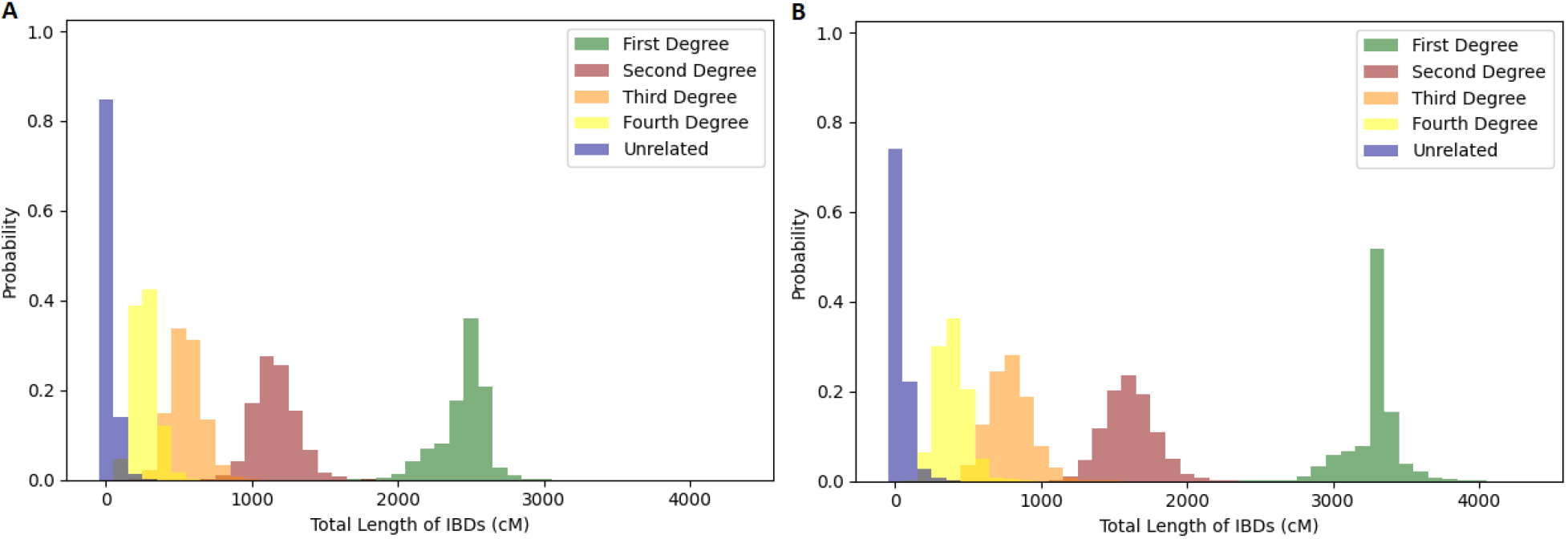
Probability Distributions of Sum of Length of IBDs on Simulated Dataset. (A) x-PBWT-Query. (B) RaPID-Query-2.0.

The fourth-degree distribution is differentiated using the sum of length of IBDs from RaPID-Query result from other distributions. Similar to the test performed on UK biobank dataset, the AUC values in Table 4 (The ROC curves are in Figure S3) between each distributions were calculated, in order to quantify the degree relatedness separation. For RaPID-Query-2.0 method, the AUC value between third- and fourth-degree relationships is 98.42%, and the AUC value between fourth-degree and unrelated relationships is 99.69%. This indicates that RaPID-Query-2.0 also has the ability to differentiate up-to-fourth-degree relationships, thanks to the output high-quality IBD segments having allowed mismatched markers and refined boundaries. The AUC values calculated from the result using x-PBWT-Query method is able to separate fourth-degree from third-degree and unrelated relationships but having lower AUC values than those calculated from RaPID-Query-2.0 method. Also, the probability distributions resulted from x-PBWT-Query method have relatively smaller means than those resulted from RaPID-Query-2.0 method (See Table S2). This means it is more confident to utilize the IBD segments identified from RaPID-Query-2.0 method than that from x-PBWT-Query method for degree of relationship separations.

**Table 4:** x-PBWT-Query versus RaPID-Query-2.0: Area Under Curve (%) of Sum of Length of IBDs on Simulated Dataset

| Method | 1st Degree<br>versus<br>2nd Degree | 2nd Degree<br>versus<br>3rd Degree | 3rd Degree<br>versus<br>4th Degree | 4th Degree<br>versus<br>Unrelated |
| --- | --- | --- | --- | --- |
| x-PBWT-Query | 100.00 | 99.85 | 97.59 | 99.24 |
| RaPID-Query-2.0 | 100.00 | 99.87 | 98.42 | 99.69 |

From the result, it is also noticed that RaPID-Query is very fast and memory-efficient for querying autosomal chromosomes of the entire UK biobank dataset. Excluding the panel preprocessing time, the average CPU time of an individual haplotype query of RaPID-Query-2.0 method is 2.76 seconds with 1.04 standard deviation. The average wall time is 3.05 seconds with 1.17 standard deviation. Naseri et al.’s PBWT-Query and L-PBWT-Query methods with 700 SNPs takes 20 seconds and 6 seconds on average per query for the UK biobank dataset (excluding the panel computing and loading time) (Naseri *et al*., 2019a). The highest peak of memory size RaPID-Query-2.0 method used to hold the pre-processed panels in memory is 1.2 terabytes, as for the PBWT-Query and L-PBWT-Query it is 2.4 terabytes and 4.7 terabytes, respectively (Naseri *et al*., 2019a). This shows RaPID-Query-2.0 method outperforms PBWT-Query and L-PBWT-Query from both the time and space perspectives. RaPID-Query has the potential power of being scalable to population-scale cohorts with feasible query time and memory usage.

## 4 Discussion

RaPID-Query has the ability of detecting high-quality IBD segments efficiently for a given individual against a population panel. It is shown that the false negative (or power) rate and the false positive (or accuracy) rate of RaPID-Query are comparable to the state-of-the-art all-vs-all IBD detection method. The query search computation time is extraordinary compared to the previous methods, as one haplotype query takes around three seconds to identify IBD segments in 22 autosomal chromosomes of entire UK biobank dataset, using the conventional genealogical search threshold.

The inferred IBD segments from RaPID-Query are available for further downstream analysis, such as relationship inference. It is shown that the IBD segments are high-grade and the sum of length of those IBDs is able to categorize at least 97% individual pairs by the relationship up to 4th degree. There is a potential that the quality relationship inference for higher degrees is reachable, if the inference method utilize not only the sum of length of IBD segments, but the combination of the number, the lengths, or the locations of IBD segments, or even demographic data as Erlich et al. used in their long-range familial search pipeline (Erlich *et al*., 2018). The multi-step IBD-segment-based relationship inference methods, for instances, CREST (Qiao *et al*., 2021), DRUID (Ramstetter *et al*., 2018), ERSA (Li *et al*., 2014), IBDkin (Zhou *et al*., 2020b), PONDEROSA (Williams *et al*., 2020), may benefit from RaPID-Query if they perform their analysis based on some high-quality IBD segments produced by RaPID-Query. If multiple individuals are needed for pedigree constructions or machine learning models during the inference, it is easy to run RaPID-Query on a parallel querying basis as each query is independent.

## Code Availability

The RaPID-Query program is available at https://github.com/ucfcbb/RaPID-Query.

## Acknowledgements

This research was conducted using the UK Biobank Resource under Application Number 24247.

