## Supplementary Material for "RaPID-Query for Fast Identity by Descent Search and Genealogical Analysis"

### 1 RaPID-Query Random Projection Example

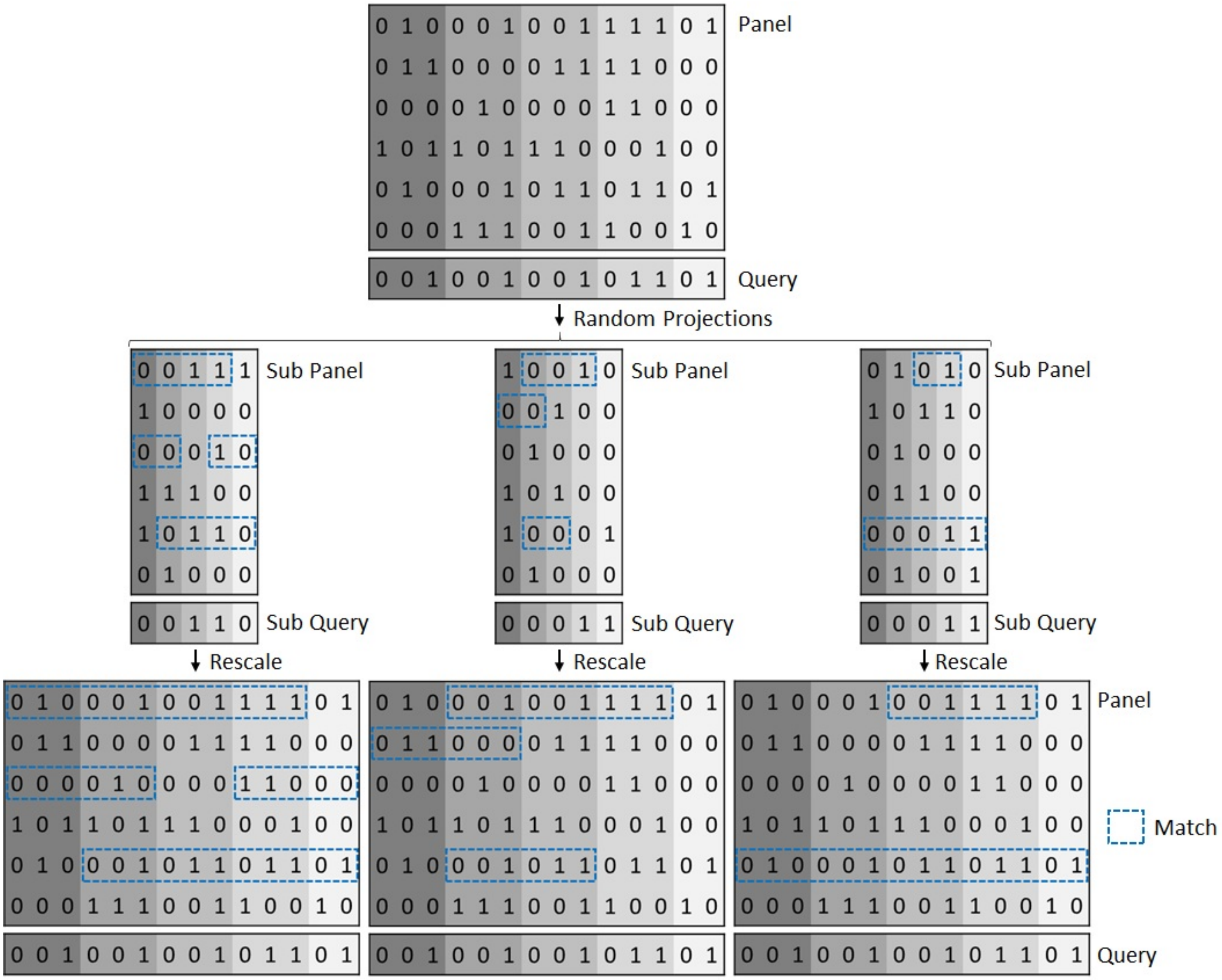


Figure S1: RaPID-Query Random Projection Example (n=14, w=3, r=3, L=6). The shading color indicates the window. One site is randomly sampled (weighted on the minor allele frequency) for each window to form $r = 3$ sub panels having $\lceil \frac{n}{w} \rceil = \lceil \frac{14}{3} \rceil = 5$ sites. The matches with at least $\frac{L}{w} = \frac{6}{3} = 2$ length in sub panels are identified by using x-PBWT-Query algorithm, and then rescaled to the original resolution.

#### 2 x-PBWT-Query Algorithm

**Algorithm S1** x-PBWT-Query

$f_L = 0, g_L = 0, e = 0, f_e = 0, g_e = m, i = 0, h =$ null
**for** $k$ from 0 to $n-1$ **do**
    // *update site index currently* $L$ *away from* $k$
    **while** $i < k + 1 - L$ **do**
        $i = i + 1$
    // *report match*
    **if** $h$ is not null **then**
        $f_r = getBlockIndicator(f_L, k, \overline{z[k]}, \textbf{true})$
        $g_r = getBlockIndicator(g_L, k, \overline{z[k]}, \textbf{false})$
        **for** $j$ from $f_r$ to $g_r - 1$ **do**
            **if** $p[k][j]$ is in $h$ **then**
                report $(p[k][j], h[p[k][j]], k-1)$
                $h.remove(p[k][j])$
    // *update search block*
    $f'_e = getBlockIndicator(f_e, k, z[k], \textbf{true})$
    $g'_e = getBlockIndicator(g_e, k, z[k], \textbf{false})$
    **if** $f'_e < g'_e$ **then**
        $e' = e$
    **else**
        **if** $f'_e == 0$ **or** $f'_e == m$ **then**
            $e' = k$
        **else**
            $e' = d[k][f'_e] - 1$
        **if** $(f'_e == m$ **and** $z[e'] == X[p[k][m-1]][e'])$ **or** $(f'_e > 0$ **and** $f'_e < m$ **and** $z[e'] == 0)$ **then**
            $f'_e = f'_e - 1$
            **while** $e' \geq 1$ **and** $z[e'-1] == X[p[k][f'_e]][e'-1]$ **do**
                $e' = e' - 1$
            **while** $f'_e > 0$ **and** $d[k][f'_e] \leq e'$ **do**
                $f'_e = f'_e - 1$
        **else if** $(g'_e == 0$ **and** $z[e'] == X[p[k][0]][e'])$ **or** $(g'_e > 0$ **and** $g'_e < m$ **and** $z[e'] == 1)$ **then**
            **while** $e' \geq 1$ **and** $z[e'-1] == X[p[k][g'_e]][e'-1]$ **do**
                $e' = e' - 1$
            $g'_e = g'_e + 1$
            **while** $g'_e < m$ **and** $d[k][g'_e] \leq e'$ **do**
                $g'_e = g'_e + 1$
        **else**
            $f'_e = 0, g'_e = m, e' = k + 1$
    $e = e', f_e = f'_e, g_e = g'_e$

**Algorithm S1** x-PBWT-Query (Continued)

```
// update match block
if f_L < g_L then
    f'_L = getBlockIndicator(f_L, k, z[k], true)
    g'_L = getBlockIndicator(g_L, k, z[k], false)
else
    f'_L = f_L
    g'_L = g_L
if f'_L == g'_L then
    if e == k + 1 − L then
        for j from f_e to g_e − 1 do
            h[p[k][j]] = e
        f'_L = f_e, g'_L = g_e
if f'_L < g'_L then
    while d[k][f'_L] ≤ k + 1 − L do
        f'_L = f'_L − 1
        h[p[k][f'_L]] = i
    while d[k][g'_L] ≤ k + 1 − L do
        h[p[k][g'_L]] = i
        g'_L = g'_L + 1
f_L = f'_L, g_L = g'_L
```

**Algorithm S2** Get Search and Match Block Indicator

```
function getBlockIndicator(indicator, k, site_value, is_indicator_f)
    if indicator == m then
        if is_indicator_f then
            if site_value == 0 then
                indicator = u[k][0]
            else
                indicator = v[k][0]
        else
            if site_value == 0 then
                indicator = v[k][0]
            else
                indicator = m
    else
        if site_value == 0 then
            indicator = u[k][indicator]
        else
            indicator = v[k][indicator]
    return indicator
```

### 3 Collect Detected Match Algorithm

**Algorithm S3** Collect Detected Match

```
for k from 0 to n − 1 do
    for each new detected match of haplotype id_new starting at h_new do
        // merge detected matches ending at k
        if h_new is not in B_a[k][id_new].H then
            B_a[k][id_new].H.add(h_new)
            B_a[k][id_new].H.h_new.count = 1
        for each h in B_a[k][id_new].H do
            if h > h_new then
                B_a[k][id_new].H.h.count + +
        // merge detected matches ending at less than k
        h_min = min_{h∈B_a[k][id_new].H}(B_a[k][id_new].H.h)
        i = k − w
        while i ≥ h_min do
            for each h in B_a[i][id_new].H do
                if h ≥ h_min then
                    B_a[i][id_new]H.h.count + +
            i = i − w
```

### 4 Identify Fragmented IBD Algorithm

**Algorithm S4** Identify Fragmented IBD

```
for k from 0 to n − 1 do
    for each id in B_a[k] do
        for each h in B_a[k][id].H do
            if B_a[k][id].H.h.count < c then
                B_a[k][id].H.remove(h)
        if B_a[k][id] is null then
            B_a[k].remove(id)
```

### 5 Consolidate Candidate IBD Algorithm

**Algorithm S5** Consolidate Candidate IBD

```
for k from 0 to n − 1 do
    for each id in B_a[k] do
        // merge overlapped fragmented IBDs ending at k
        h_min = min_{h∈B_a[k][id].H}(B_a[k][id].H.h)
        for each h in B_a[k][id].H do
            if B_a[k][id].H.h > h_min then
                B_a[k][id].H.remove(h)
        // merge overlapped fragmented IBDs ending at less than k
        i = k − w
        while i ≥ h_min − 1 do
            if id is in B_a[i] then
                h_prev = B_a[i][id].H.h
                B_a[i][id].H.remove(h)
                B_a[i].remove(id)
                if h_prev < h_min then
                    B_a[k][id].H.h = h_prev
                    break
            i = i − w
```

### 6 Refine Candidate IBD Algorithm

**Algorithm S6** Convert Candidate IBD Form

```
for k from n − 1 to 0 do
    for each id in B_a[k] do
        B_m[id].S.push((B_a[k][id].H.h, k))
```

**Algorithm S7** Refine Candidate IBD

**for** each $(id, [h_{full}, t_{full}])$ candidate IBD from querying full panel **do**
  **if** $B_m[id]$ is not null **then**
    $(h_{low}, t_{low}) = B_m[id].S.top()$
    **while** $t_{low} < t_{full}$ **do**
      $B_m[id].S.pop()$
      $(h_{low}, t_{low}) = B_m[id].S.top()$
    **if** $(h_{low}, t_{low})$ has overlap with $(h_{full}, t_{full})$ **then**
      **if** $B_m[id].R$ is null **then**
        **if** $t_{low} \leq t_{full}$ **then**
          $B_m[id].S.pop()$
          report $(id, [max(h_{low}, h_{full}), t_{low}])$
        **else**
          $B_m[id].R = (max(h_{low}, h_{full}), t_{full})$
      **else**
        $(h_R, t_R) = B_m[id].R$
        **if** $t_{low} \leq t_{full}$ **then**
          $B_m[id].S.pop()$
          **if** $h_{full} - t_R \leq g_{max}$ **then**
            report $(id, [h_R, t_{low}])$
          **else**
            report $(id, [h_R, t_R])$
            report $(id, [h_{full}, t_{low}])$
          $B_m[id].R = \text{null}$
        **else**
          **if** $h_{full} - t_R \leq g_{max}$ **then**
            $B_m[id].R = (h_R, t_{Full})$
          **else**
            report $(id, [h_R, t_R])$
            $B_m[id].R = (h_{full}, t_{Full})$

### 7 Probability Distributions of Sum of Length of IBDs on UK Biobank Dataset

| **Degree Distribution** | **x-PBWT-Query** | | **RaPID-Query-2.0** | |
|---|---|---|---|---|
| | Mean | Standard Deviation | Mean | Standard Deviation |
| 1st Degree | 2594.94 | 754.78 | 3269.66 | 544.51 |
| 2nd Degree | 1287.84 | 347.56 | 1591.89 | 330.75 |
| 3rd Degree | 591.80 | 195.69 | 756.28 | 158.20 |
| Unrelated | 0.22 | 5.78 | 0.23 | 6.17 |

Table S1: Probability Distributions of Sum of Length of IBDs Parameters on UK Biobank Dataset

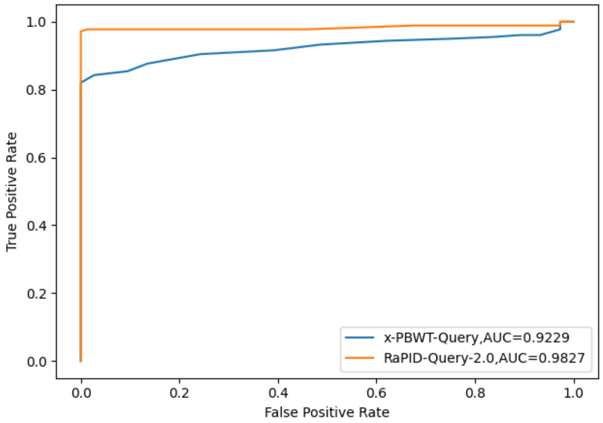


(a) 1st Degree vs 2nd Degree

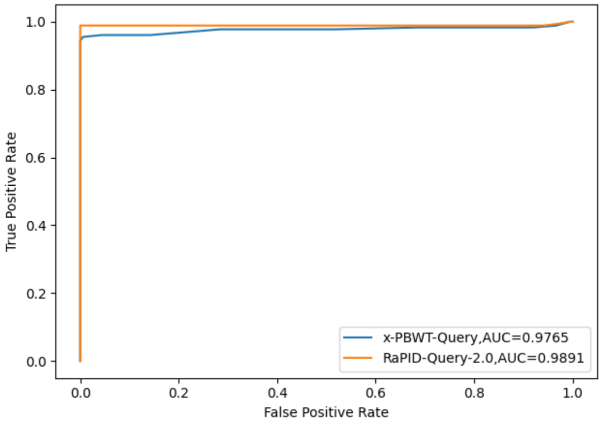


(b) 1st Degree vs 3rd Degree

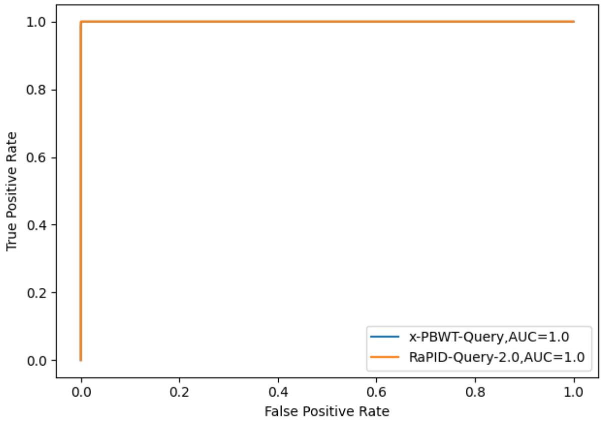


(c) 1st Degree vs Unrelated

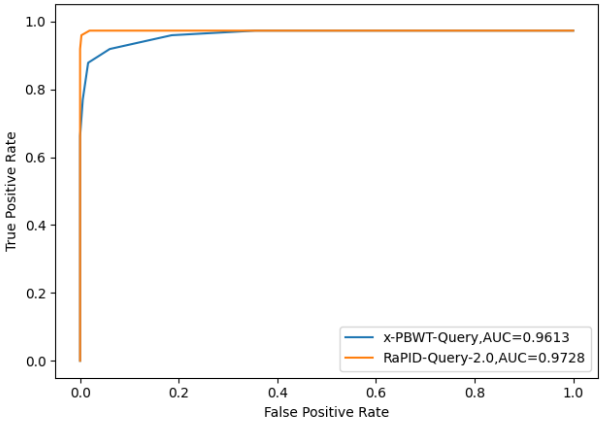


(d) 2nd Degree vs 3rd Degree

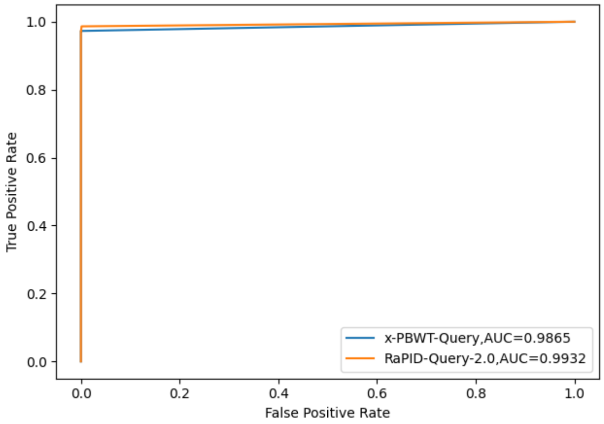


(e) 2nd Degree vs Unrelated

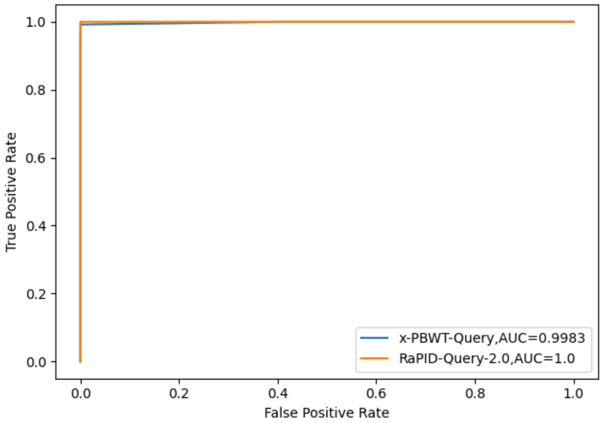


(f) 3rd Degree vs Unrelated

Figure S2: ROC Curves and AUC Values of Sum of Length of IBDs on UK Biobank Dataset

### 8 Probability Distributions of Sum of Length of IBDs on Simulated Dataset

| **Degree Distribution** | **x-PBWT-Query** | | **RaPID-Query-2.0** | |
|---|---|---|---|---|
| | Mean | Standard Deviation | Mean | Standard Deviation |
| 1st Degree | 2448.88 | 163.15 | 3276.03 | 167.17 |
| 2nd Degree | 1158.86 | 145.00 | 1605.77 | 178.79 |
| 3rd Degree | 553.29 | 115.86 | 788.05 | 148.96 |
| 4th Degree | 267.64 | 82.88 | 391.27 | 107.71 |
| Unrelated | 16.74 | 41.34 | 30.21 | 56.36 |

Table S2: Probability Distributions of Sum of Length of IBDs Parameters on Simulated Dataset

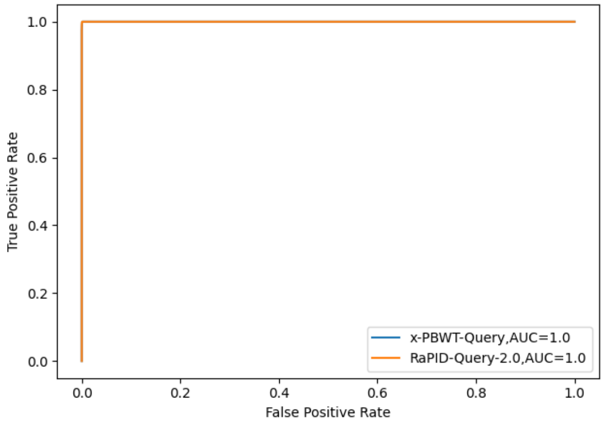


(a) 1st Degree vs 2nd Degree

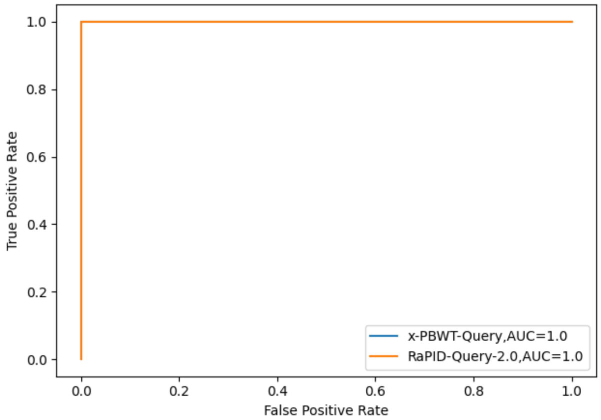


(b) 1st Degree vs 3rd Degree

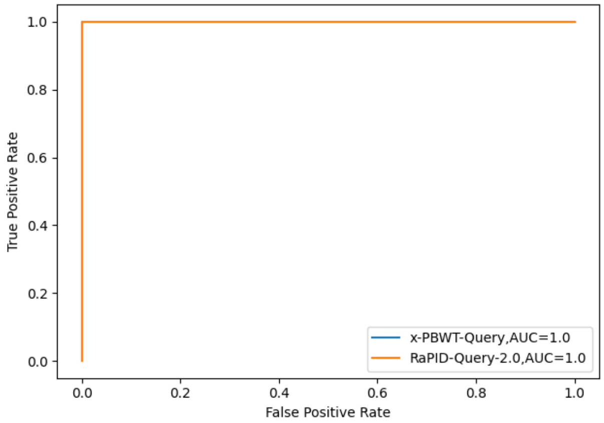


(c) 1st Degree vs 4th Degree

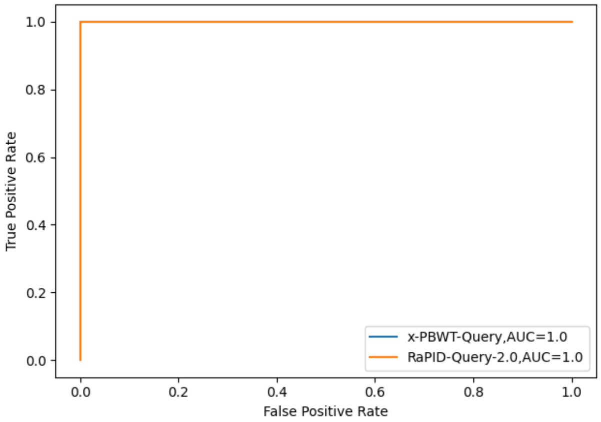


(d) 1st Degree vs Unrelated

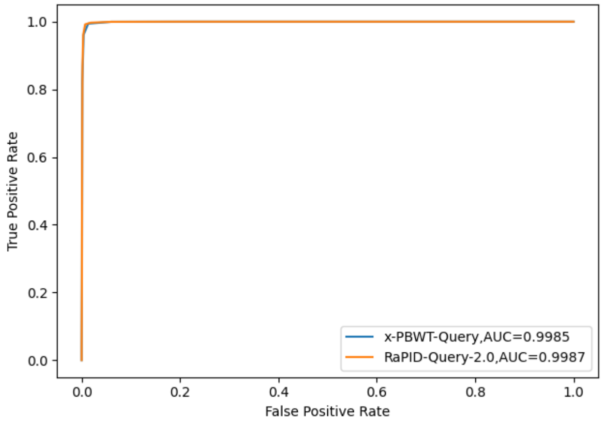


(e) 2nd Degree vs 3rd Degree

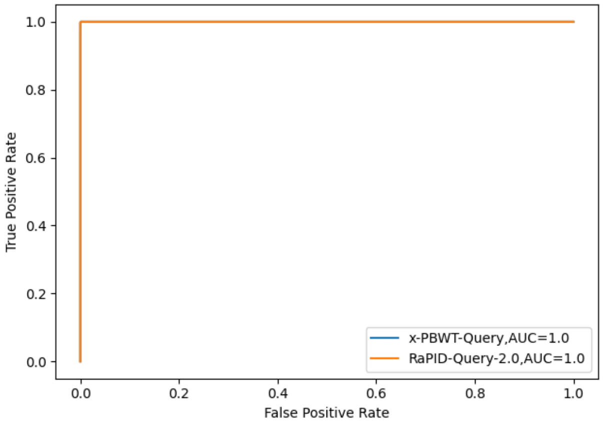


(f) 2nd Degree vs 4th Degree

Figure S3: ROC Curves and AUC Values of Sum of Length of IBDs on Simulated Dataset

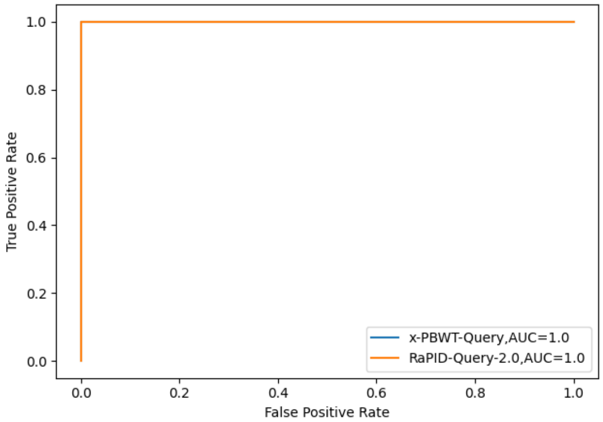


(a) 2nd Degree vs Unrelated

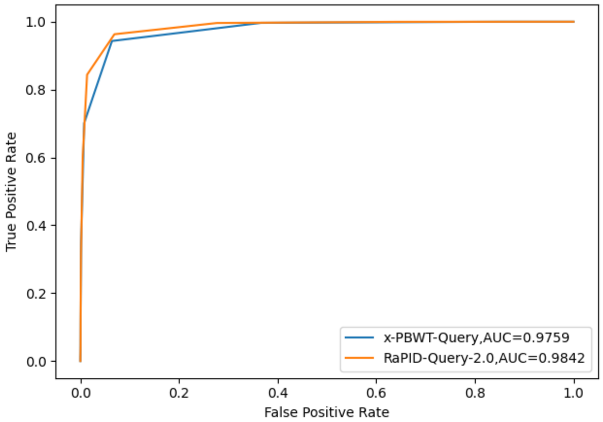


(b) 3rd Degree vs 4th Degree

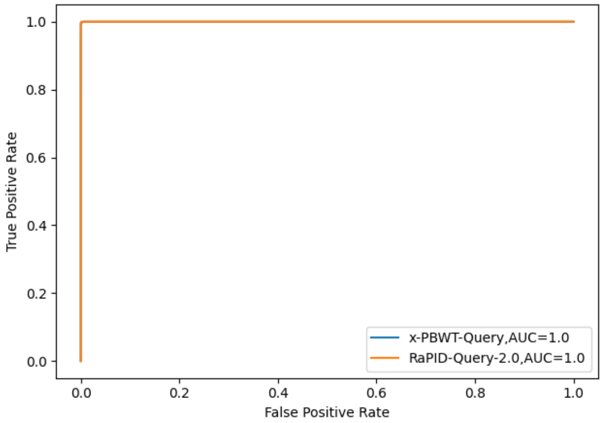


(c) 3rd Degree vs Unrelated

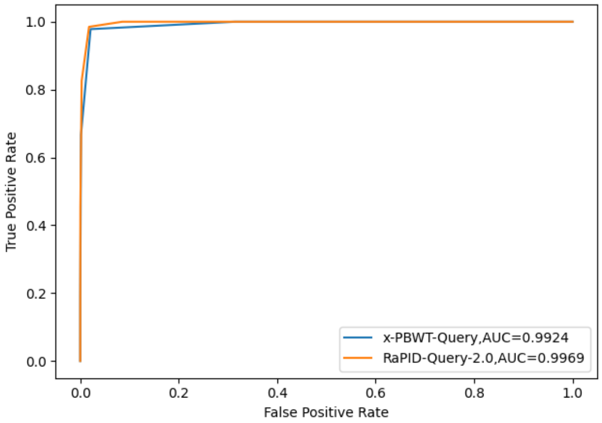


(d) 4th Degree vs Unrelated

Figure S3: ROC Curves and AUC Values of Sum of Length of IBDs on Simulated Dataset (Continued)
